# Mutation of a single cysteine in CaMKIIδ protects the heart from ischemia-reperfusion Injury

**DOI:** 10.64898/2026.04.27.721066

**Authors:** Nathália Rocco-Machado, Junhui Sun, Audrey Noguchi, Danielle A. Springer, Chengyu Liu, Elizabeth Murphy, Rodney L. Levine

## Abstract

CaMKIIδ is the dominant isozyme of Ca^2+^/calmodulin-dependent protein kinase II in the heart. Under certain pathological conditions, it can be oxidized, causing a constitutive activation that can lead to cardiac failure. We recently showed that, in purified CaMKIIδ exposed to oxidative conditions, a disulfide link formed between Cys273 and Cys290 causes this autonomous activation. Cys273 has a low pKa that facilitates the oxidation of its thiol to a sulfenic acid at physiological pH. Does this matter *in vivo*? To answer that question, we created a transgenic mouse with Cys273 mutated to serine (CaMKIIδ^C273S^) to prevent disulfide formation. We conducted a detailed assessment of cardiac function at rest and in a dobutamine stress test. We found that the CaMKIIδ Cys273Ser mutation does not have deleterious effects on cardiac physiology. Then, we assessed whether the mutation would protect the heart from ischemia-reperfusion in the Langendorff model. The CaMKIIδ^C273S^ mouse had improved cardiac function and decreased infarct size compared to the wild-type mouse. We conclude that blocking disulfide formation at Cys273 protects the heart against ischemia-reperfusion injury. Drugs that specifically target Cys 273 may be therapeutic in human cardiac disease.

## Introduction

Myocardial ischemia-reperfusion injury occurs when the blood supply to the heart is interrupted (ischemia) and then re-established (reperfusion), leading to calcium overload and an increase in reactive oxygen species (ROS) (1-3). Ischemia-reperfusion injury can induce cardiomyocyte death and result in infarction (1-3). Currently, there is no effective therapy to prevent or reduce it (3, 4). The combination of intracellular calcium and ROS increase can lead to the serine/threonine kinase Ca^2+^/calmodulin-dependent protein kinase II (CaMKII) oxidation and overactivation (5-8). CaMKII overactivation during reperfusion has been causally linked to early ischemia-reperfusion arrhythmias, heart dysfunction, and cell death (5-7, 9, 10).

CaMKII has four isoforms, α, δ, β, and γ, encoded by different genes. CaMKIIδ is the dominant form in the heart and, in states of increased physiological demand, it can regulate pathways that influence, for example, intracellular Ca^2+^ regulation and excitation-contraction coupling (11, 12), and gene transcription (13, 14). However, under pathological conditions, it can be overactivated and lead to cardiac failure (5-7, 9, 10). CaMKIIδ oxidation promotes sustained, autonomous kinase activity (Ca^2+^/calmodulin independent activity) (5, 8).

The CaMKIIδ oxidation site was initially believed to be methionine residues 281 and 282, and an antibody specific for oxidized CAMKIIδ was thought to recognize the oxidized methionine residues (5). However, our recent publication unequivocally established that in purified, recombinant CaMKIIδ exposed to oxidative conditions, it is a disulfide link formed between Cys273 and Cys290 that confers CaMKIIδ autonomous activation (8). Methionine residues are not oxidized, and the antibody specifically recognizes the disulfide-linked Cys273. Cys273 has a low apparent pKa that facilitates the oxidation of its thiol to sulfenic acid at physiological pH (8). The reactive sulfenic acid then attacks the thiol of Cys290. Does this matter *in vivo*? To answer that question, we created a knockin mouse line with Cys273 mutated to serine (CaMKIIδ^C273S^) so that the disulfide bond cannot form. In our previous publication (8), we showed that mutation of CaMKIIδ Cys273 to serine has similar Ca^+2^/calmodulin-dependent activity when compared to the WT enzyme, however, it does not present autonomous activation induced by oxidation. We recognized that the mutant CaMKIIδ could have deleterious effects on cardiac physiology, so we conducted a detailed assessment of cardiac function at rest and in a dobutamine stress test. To determine whether the mutation protected against ischemia-reperfusion injury, we utilized the Langendorff model of cardiac ischemia-reperfusion. We show that cardiac physiology is unaffected by the introduction of the mutation, while the heart is significantly protected from ischemia-reperfusion injury.

## Material and methods

### Generation of the CaMKIIδ^C273S^ knockin mouse line

All mice described in this work were generated on C57BL/6 background. Mice were treated in accordance with the Guide for the Care and Use of Laboratory Animals (15), and the study was approved by the Animal Care and Use Committee of the National Heart, Lung, and Blood Institute (H-0120R6). We obtained informed written consent. The study protocol requires humane endpoints, but no mice required such euthanasia, and none died before meeting criteria. The criteria for euthanasia were: tumors greater than 2 cm, ulcerated, or interfering with eating or drinking; lethargy; dehydration; reluctance to move about the cage; non-responsiveness to touch; or decreased food/water consumption. No randomization was used to allocate experimental units to control and treatment groups, and no strategy was used to minimize potential confounders.

Mice were housed in ventilated cage racks in an Association for Assessment and Accreditation of Laboratory Animal Care International (AAALAC) accredited animal facility at the National Institutes of Health (NIH), Bethesda, MD, with ad lib food and water. The animal facility housing room was maintained on a 12 h light: dark schedule, and room temperature and humidity followed the Guide. NIH Division of Veterinary Resources provided daily health checks and health care to the animals. Details of the dobutamine stress test and of the ischemia-reperfusion are described below. Eleven or 12 animals with wild-type or CaMKIIδ^C273S^ genotype were utilized, a number established in previous studies to provide acceptable power in both the stress test and the ischemia-reperfusion studies.

The CaMKIIδ^C273S^ knockin mouse line was generated using CRISPR/Cas9 technology (16). Briefly, a single guide RNA (sgRNA: AGTGACTTACACAGATCCAT) was purchased from Synthego, and a single-strand oligo donor template was ordered from IDT with this sequence: (GCCAAAGACCTCATCAACAAAATGCTGACCATCAACCCTGCCAAACGTATCACAG CCTCTGAGGC**A**CTGAAACACCCATGGATC**TCC**GTAAGTCACTTTCGCATGCACCCC AGGAGCCAAACTGAAGGATAAACCCCTAGTCAC). The bold and underlined codon encodes the desired Ser residue. The single bold nucleotide (A) is a synonymous mutation in the PAM of the sgRNA, introduced for the purpose of stopping Cas9 after the donor template has been successfully knocked in. The sgRNA (20 ng/μL) and donor oligos (100 ng/μL) were co-microinjected with Cas9 mRNA (50 ng/ μL, TriLink BioTechnologies) into the cytoplasm of zygotes collected from C57BL6/ mice. Injected embryos were cultured overnight in KSOM medium (Millipore) in a 37°C incubator with 6% CO_2_. The next morning, embryos that had reached the 2-cell stage of development were implanted into the oviducts of pseudo-pregnant surrogate mothers (CD1 mice, Charles River Laboratory). Offspring born to the foster mothers were genotyped using PCR followed by Sanger DNA sequencing to identify mice with the desired nucleotide changes.

While it is still debatable whether an off-target effect of this technology is a major concern, whole genome sequencing studies have shown that off-target mutations are rare in germline-mutated mice (17, 18). In addition, a large scale whole genome sequencing of 36 mice with 60x coverage also found that off-target mutations are rare (19), and the mice in this study were generated in the same core facility with the same experimental conditions. Specifically, we injected the Cas9 mRNA and guide RNAs into zygotes, in which they should be stable for only a few days. Other off-target studies used somatic cell lines that constitutively expressed Cas9 and guide RNAs. In addition, the original founder mice had been bred with unedited mice for several generations during CaMKIIδ^C273S^ knockin mouse line establishment and colony expansion. Therefore, rare off-target mutations, if any, were likely diluted out before the mice were used for experiments.

In all the experiments conducted here, we compared WT mice with CaMKIIδ^C273S^ homozygous mice, both obtained by mating two CaMKIIδ^C273S^ heterozygous mice.

### Dobutamine Cardiac Stress Test

Mice of both genders, age 2-5 months, were lightly anesthetized with 5% isoflurane in an induction chamber and then placed supine on a heated platform with ECG leads, a rectal temperature probe, and a nose cone at 2% isoflurane to perform echocardiography exams before and after administering constant rate dobutamine infusions. Heart images were acquired using the Vevo2100 ultrasound system (VisualSonics, Toronto, Canada) with a 30 MHz ultrasound probe (VisualSonics, MS-400 transducer). After the baseline scan, the mice received constant rate infusions of dobutamine (0.625 mg/ml in normal saline containing 5% dextrose) via the tail vein using an infusion syringe pump (Harvard Apparatus). The low dose infusion rate was 10 μg/kg/min. After the heart rate reached a steady state as determined by the ECG, the rate was increased to 40 μg/kg/min, and the scans were repeated.

### Ischemia-reperfusion model

Mice of both genders, aged between 2-5 months old, were studied. After anesthesia with pentobarbital (50-70 mg/kg body weight) and anticoagulation with heparin (50 USP units per 25 g mouse). When anesthetized mice lost the toe pinch reflex, a thoracotomy was performed, and the heart was quickly excised and placed in ice-cold Krebs-Henseleit buffer (in mmol/l: 120 NaCl, 11 D-glucose, 25 NaHCO_3_, 1.75 CaCl_2_, 4.7 KCl, 1.2 MgSO_4_, and 1.2 KH_2_PO_4_). The aorta was cannulated, and the heart was perfused with Krebs-Henseleit buffer (oxygenated with 95% O2/5% CO_2_ and maintained at pH 7.4) in retrograde fashion at a constant pressure of 100 cm of water at 37°C. A latex balloon connected to a pressure transducer was inserted into the left ventricle of the perfused heart to monitor left ventricular developed pressure (LVDP). LVDP was recorded and digitized using a PowerLab system (ADInstruments, Colorado Springs, CO). After an equilibration period of 20 minutes, hearts were subjected to 20 minutes of no-flow ischemia followed by 120 minutes of reperfusion. The rate-pressure product (RPP=LVDP x heart rate) was used as an index of cardiac contractile function. Post-ischemic functional recovery was expressed as a percentage of the pre-ischemic RPP during the equilibration period. At the end of the 120 min of reperfusion, hearts were perfused with 1% (wt/vol) of 2,3,5-triphenyltetrazolium chloride (TTC) and incubated in TTC at 37°C for 15 min, followed by fixation in 10% (wt/vol) formaldehyde. Infarct size was expressed as the percentage of the total cross-sectional area of the ventricles.

### CaMKII expression in the heart

The hearts from 8 different mice of both genders, age 2-5 months (4 WT and 4 CaMKIIδ^C273S^), were dissected and snap frozen in liquid nitrogen for protein analysis. Protein lysates were prepared with 50mM Tris-HCl pH 7.5, 1mM phenylmethylsulfonyl fluoride (PMSF) (Sigma 10837091001), 3% SDS, and supplemented with 1x protease inhibitor (Sigma P2714-1BTL). Lysates were then incubated on ice for 15 min and centrifuged for 15 min at 15000 rcf at 4°C. SDS-page and Western blot were performed as described below.

### CaMKIIδ expression and half-life determination in HEK293 cells

HEK293 cells were cultured in Dulbecco’s modified Eagle’s medium (Invitrogen 11965092) with 10% fetal calf serum, and 1% penicillin and streptomycin at 37 °C in a humidified atmosphere of 5% CO_2_ and 95% air. Cells were transfected with wild-type or C273S CaMKIIδ mutant, using calcium phosphate (Invitrogen K278001). Forty-eight hours after transfection, cells were treated with 50 μg/ml cycloheximide (CHX) (Sigma 239763). At 0, 1, 2, and 3 hours after incubation, cells were collected and lysed with 25 mM Tris–HCl, pH 7.4, 150 mM NaCl, 1 mM EDTA, 1% Nonidet P-40, 5% glycerol, 1 mM PMSF (Sigma 10837091001), and 1× protease inhibitor cocktail (Sigma P2714-1BTL). Lysates were cleared by centrifugation for 20 min at 20,800g at 4 °C. Western blot analysis was performed as described below.

### SDS-page and Western blot

HEK293 supernatant or heart lysate supernatant protein concentration was determined by the BCA method (Thermo Fisher Scientific P6071161). Protein samples were mixed with 2× Novex Tris-glycine SDS sample buffer (Thermo Fisher Scientific LC2676) supplemented with 1 mM 2-mercaptoethanol and heated for 3 min at 100°C when using the primary antibody recognizing CaMKII delta (Invitrogen PA5-22168-1:10,000). 20 μg (HEK293) or 50ug (Heart) of protein was loaded onto each lane, and SDS-PAGE was performed on Novex 10 to 20% Tris-glycine gel (Thermo Fisher Scientific XP10200BOX) in Novex Tris-glycine SDS running buffer (Invitrogen LC2675) at a maximum 125 V for 2 h. Proteins were transferred to nitrocellulose membrane using the Trans-Blot Turbo Mini 0.2 µm Nitrocellulose Transfer (BioRad 1704158) at 25 V, 1.3–2.5 A, 7 minutes using the BioRad Trans-Blot Turbo system. Membranes were blocked for 30 min with blocking buffer (Li-Cor 927-70001), followed by incubation in blocking buffer supplemented with 0.05% Tween 20 and 1:10,000 diluted primary antibodies overnight on a rocking platform at 4 C. The next day, incubation was continued for 1 more hour at room temperature and then washed with 1× PBS, pH 7.4, twice for 10 min, followed by incubation in blocking buffer (0.05% Tween 20) supplemented with a 1:25,000 dilution of LICOR IRDye 800CW Goat anti-Rabbit IgG Secondary Antibody (LICOR 92632211) for 20 min at room temperature on a rocking platform. Membranes were then washed with PBS for 15 min with a change of buffer every 5 min. Quantitative Western blot images were obtained by scanning the membrane on a Li-Cor Odyssey infrared imaging system according to the manufacturer’s instructions. Protein loading was normalized by total protein staining (LICOR 92611016).

### Statistical Analysis

All data were analyzed with GraphPad Prism 10 software. A two-tailed Student’s t-test was used to determine the statistical significance of differences between groups, with p values <0.05 considered significant. For the parameters analyzed in Table 1, we used the Bonferroni method to adjust the statistical significance level based on the number of tests performed so that a difference between the groups was only considered significant if p < 0.008. All reported experiments were repeated independently at least three times.

**Table 1.** Baseline comparison of wild-type and CaMKIIδ^C273S^ mice and Cardiac parameters during the dobutamine stress test. **(A) Baseline comparison of wild-type and CaMKIIδ^C273S^ mice.** The values are the mean ± SE. wild-type n=11, CaMKIIδ^C273S^ n=12. P value was obtained with the unpaired two-tailed t-test. **(B) Cardiac parameters during the dobutamine** (**40 µg/kg/min) stress test**. The values are the mean ± SE. wild-type n=11, CaMKIIδ^C273S^ n=12. The p value was obtained with the unpaired two-tailed t-test. We used the Bonferroni method to adjust the statistical significance level based on the number of tests performed so that a difference between the groups was only considered significant if p < 0.008. Similarly, no significant differences were observed with the low dose dobutamine stress test (10 µg/kg/min).

| Parameter | Wild-type | C273S | P value |
| --- | --- | --- | --- |
| Animal Weight (g) | 25 ± 5.9 | 26 ± 4.3 | 0.77 |
| Heart Rate (beats/min) | 504 ± 50.1 | 513 ± 56.3 | 0.67 |
| Cardiac Output (mL/min) | 23 ± 4.8 | 23 ± 2.6 | 0.76 |
| Fractional Shortening (%) | 29 ± 1.9 | 27 ± 1.8 | 0.14 |
| Diam LV Chamber in Systole (mm) | 3 ± 0 | 3 ± 0 | 0.54 |
| Diam LV Chamber in Diastole (mm) | 4 ± 0 | 4 ± 0 | 0.86 |

**Table 1. (A) Baseline comparison of wild-type and CaMKII $\delta^{C273S}$ mice.** The values are the mean ± SE. wild-type n=11, CaMKII $\delta^{C273S}$ n=12. P value was obtained with the unpaired two-tailed t-test. **(B) Cardiac parameters during the dobutamine (40**
| Parameter | Wild-type | C273S | P value |
| --- | --- | --- | --- |
| Cardiac Output (mL/min) | 28 ± 4.7 | 27 ± 4.1 | 0.38 |
| Fractional Shortening (%) | 46 ± 4.1 | 42 ± 2.6 | 0.016 |
| Volume of LV in Systole (mm<sup>3</sup>) | 14 ± 3.9 | 16 ± 3.3 | 0.14 |
| Volume of LV in Diastole (mm<sup>3</sup>) | 62 ± 11 | 62 ± 7.7 | 0.87 |
| Diam LV Chamber in Systole (mm) | 2 ± 0 | 2 ± 0 | 0.13 |
| Diam LV Chamber in Diastole (mm) | 4 ± 0 | 4 ± 0 | 0.93 |

## Results and Discussion

### Cardiac physiology at rest and under stress is not affected by the C273S mutation

To prevent CaMKIIδ autonomous activation by disulfide formation, we used CRISPR/cas9 to generate a mouse line in which cysteine 273 was mutated to serine (Fig. 1A). The mutation did not affect CaMKII expression in the heart (Supplementary Fig.1). We then assessed the basal cardiac function of CaMKIIδ^C273S^ and wild-type mice by echocardiography. No differences were observed between the two groups, including cardiac output. (Fig. 1B-D and Table 1A).

**Figure 1.**
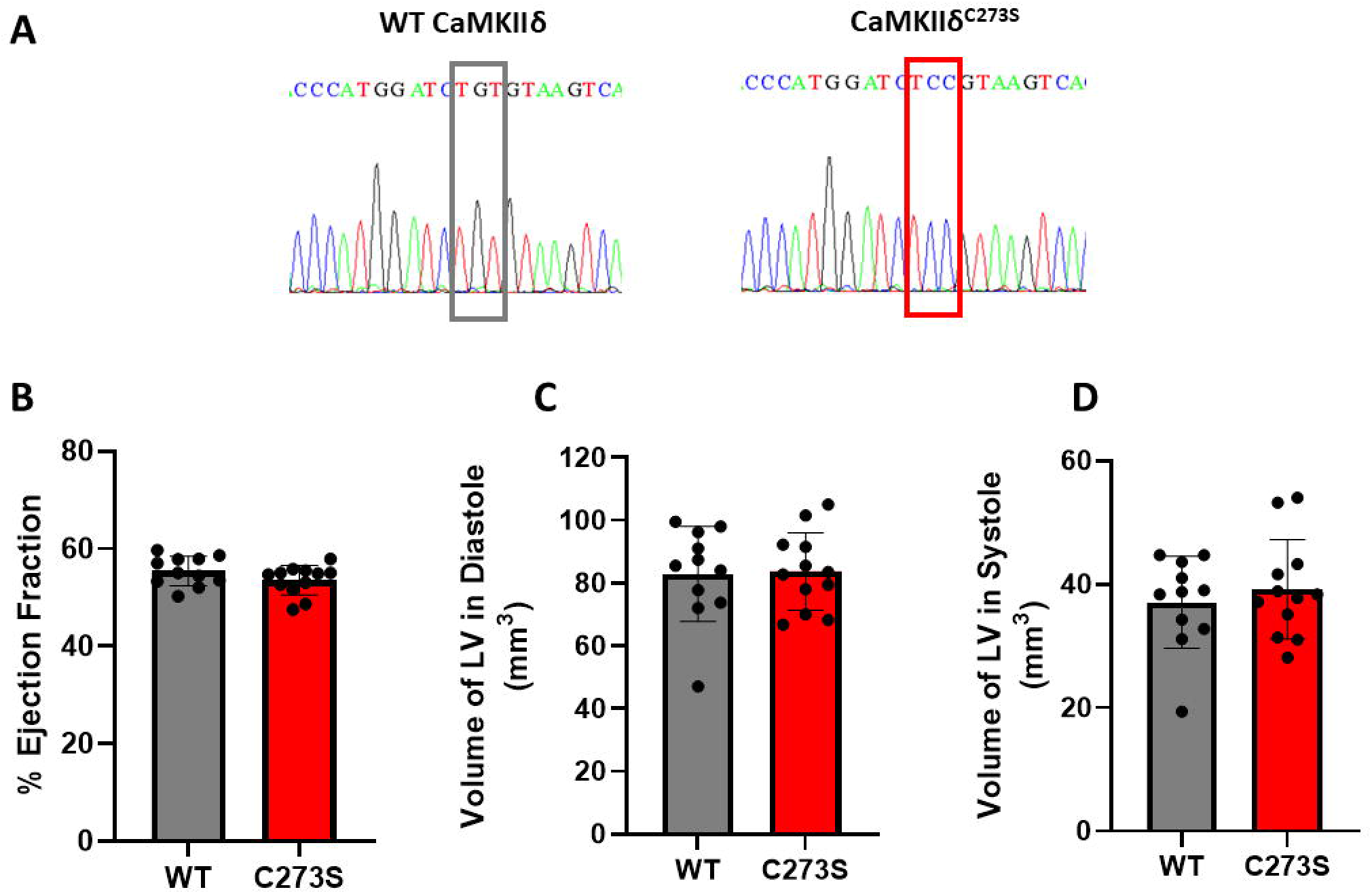
CaMKIIδ^C273S^ and wild-type mice echocardiogram under basal conditions. **(A)** The CaMKIIδ^C273S^ mouse genotype was confirmed by sequencing of tail tip DNA samples. **(B)** Wild-type (grey) and CaMKIIδ^C273S^ (red) ejection fraction. **(C)** Wild-type (grey) and CaMKIIδ^C273S^ (red) left ventricular diastolic volume. **(D)** Wild-type (grey) and CaMKIIδ^C273S^ (red) left ventricular systolic volume. Wild-type n=11, CaMKIIδ^C273S^ n=12. The difference between the groups was statistically non-significant with the unpaired two-tailed t-test.

To further characterize the cardiac function of the CaMKIIδ^C273S^ mouse, we performed dobutamine stress tests (Fig. 2 and Table 1B). Dobutamine is a β-adrenergic agonist with a high affinity for the heart receptor (β1) (20). It increases cardiac contractility and heart rate, mimicking the effects of a human exercise stress test (20). Both wild-type and CaMKIIδ^C273S^ mice responded the same way to both low and high dobutamine doses (Table 1B). Dobutamine significantly increased the ejection fraction as well as heart rate (Fig. 2A and 2B).

**Figure 2.**
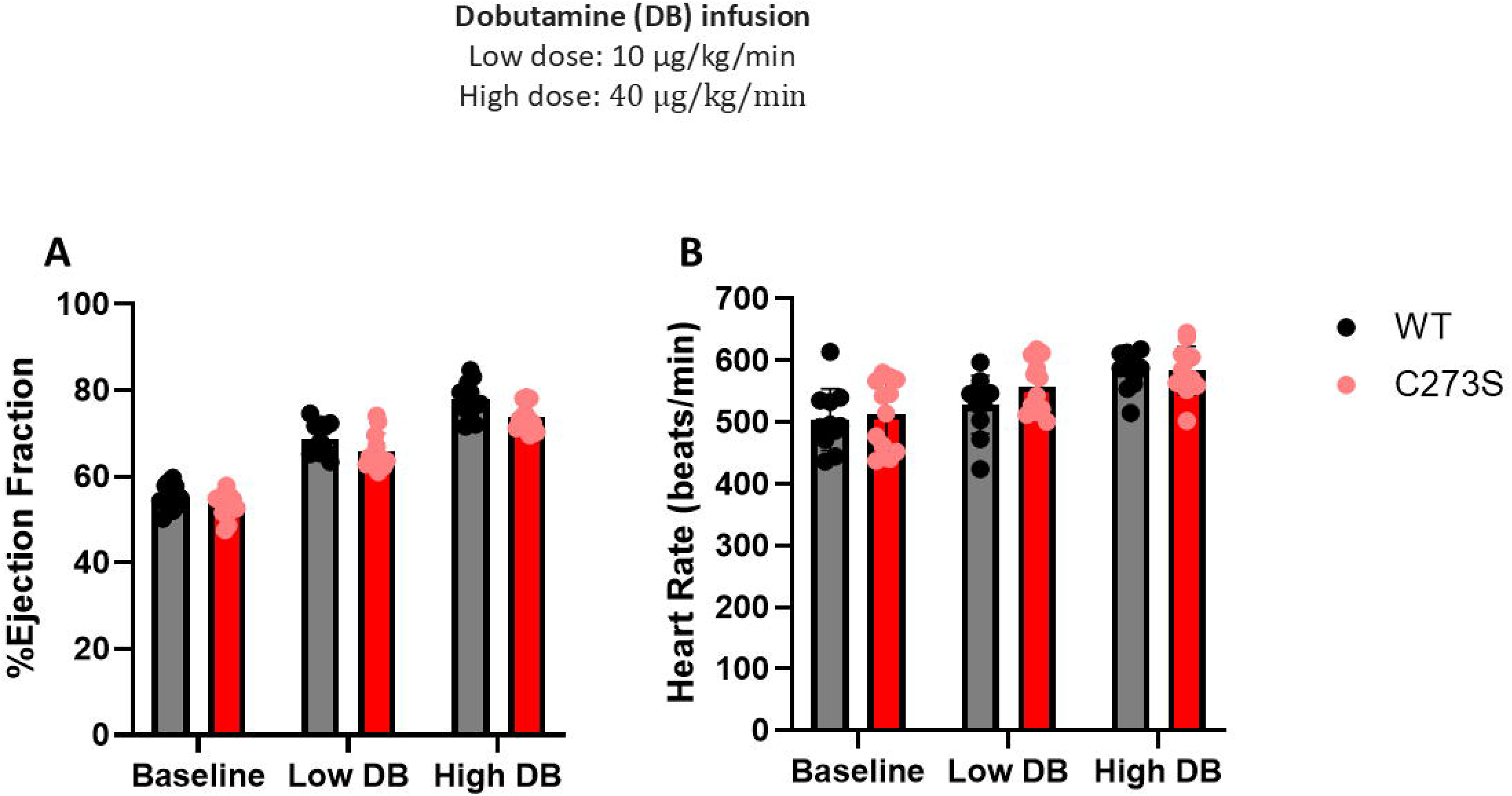
CaMKIIδ^C273S^ and wild-type mice dobutamine stress test. **(A)** Wild-type (grey) and CaMKIIδ^C273S^ (red) ejection fraction at baseline, low dobutamine dose (10 µg/kg/min), or high dobutamine dose (40 µg/kg/min). **(B)** Wild-type (grey) and CaMKIIδ^C273S^ (red) heartbeat rate at baseline, low dobutamine dose (10 µg/kg/min), or high dobutamine dose (40 µg/kg/min). Wild-type n=11, CaMKIIδ^C273S^ n=12. The comparison between wild-type and CaMKIIδ^C273S^ was statistically non-significant with the unpaired two-tailed t-test. Both wild-type and CaMKIIδ^C273S^ ejection fraction and heartbeat were statistically significantly higher after the high dobutamine dose with the unpaired two-tailed t-test.

We conclude that mutation of Cys273 to serine has no effect on cardiac physiology at rest or under stress. There were no significant differences in ejection fraction (Fig. 1B), left ventricle diastolic volume (Fig. 1C), systolic volume (Fig. 1D), nor fractional shortening (Table 1A). We note that Power and colleagues recently reported that their C273S CaMKIIδ mice (of similar age to ours) had a slight increase in left ventricle diastolic and systolic volumes and a modest reduction of ejection fraction compared to wild-type mice (21). A long-term assessment of cardiac physiology will be necessary to evaluate whether the CaMKII^δC273S^ knockin mouse line is differentially affected by aging.

### The C273S mutation protects the heart from ischemia-reperfusion injury in a Langendorff model

During the reperfusion phase of ischemia/reperfusion in the heart, angiotensin II can activate NADPH oxidase 2, increasing ROS production and leading to oxidation of CaMKII (5, 6, 10). Elevated ox-CaMKII has been shown to phosphorylate RyR2, a major calcium release channel in heart muscle cells, increasing Ca^2+^ leak and promoting atrial fibrillation and delayed afterdepolarizations (DADs) through activation of the Na^+^/Ca^2+^ exchanger (NCX). DADs can lead to arrhythmias and cardiac failure (10, 22, 23).

We hypothesized that the Cys273 mutation would be protective against ischemia-reperfusion damage. To determine whether this was the case, we compared wild-type and mutant hearts in the Langendorff model of ischemia-reperfusion (Fig. 3A). The Cys273S mouse had improved cardiac function and decreased infarct size compared to the wild-type mouse (Fig. 3B and 3C).

**Figure 3.**
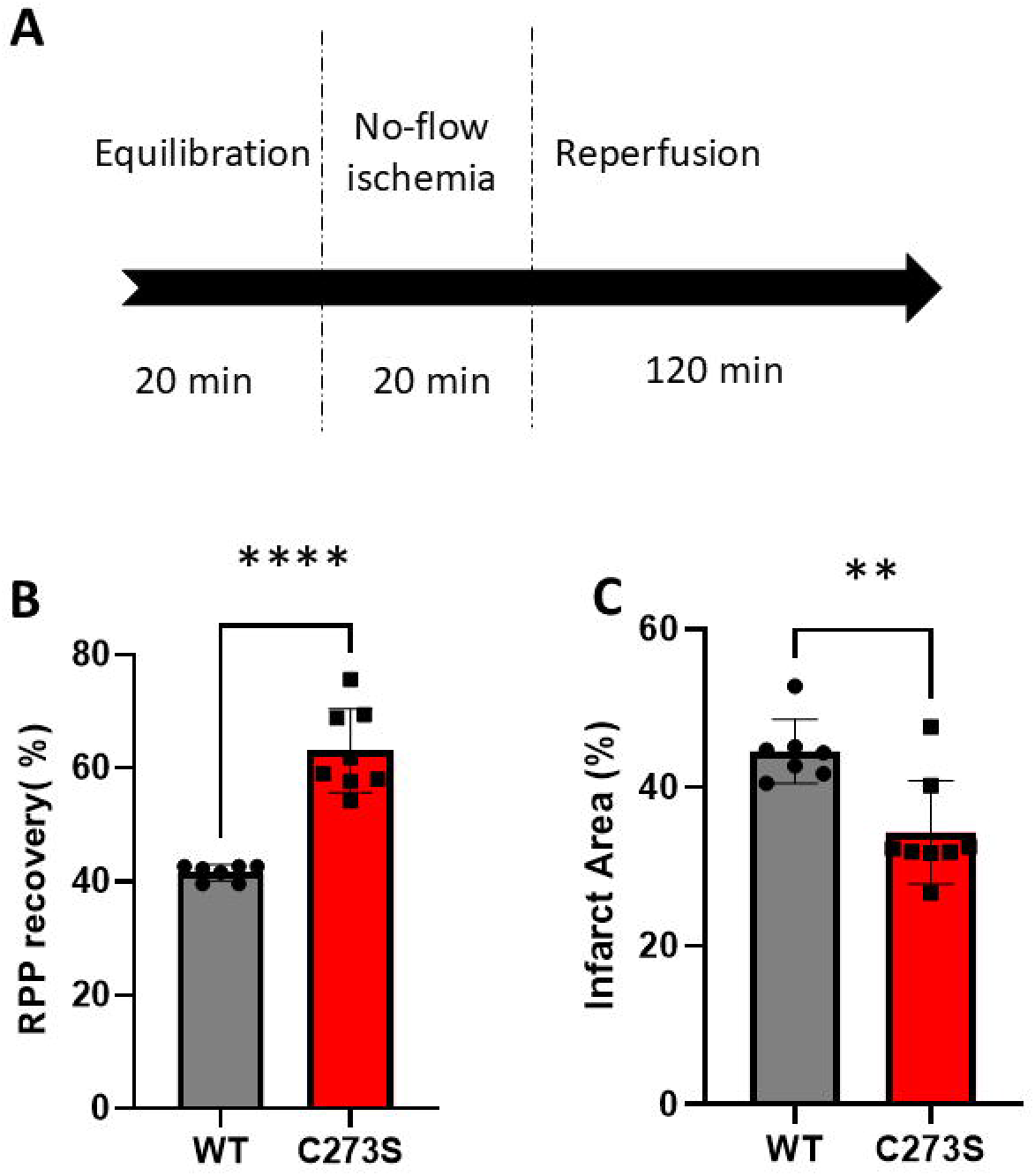
Postischemic left ventricular rate pressure (RPP) functional recovery and infarct size. **(A)** Scheme showing ischemia-reperfusion process in the Langendorff perfused heart model. **(B)** Wild-type (grey) and CaMKIIδ^C273S^ (red) postischemic left ventricular rate times pressure (RPP) as a functional measure of recovery. **(C)** Wild-type (grey) and CaMKIIδ^C273S^ (red) infarct area. Wild-type n=7, CaMKIIδ^C273S^ n=8. ****, p <0.0001. **, p = 0.003 with the unpaired two-tailed t-test.

Recently, Hansen and colleagues (22) showed that CaMKII and ROS contribute to early ischemia/reperfusion arrhythmias. They used Langendorff-perfused hearts submitted to 20 minutes of ischemia followed by 5 minutes of reperfusion. The investigators observed that when hearts were treated with the CaMKII inhibitor KN93 or the antioxidant *N*-acetylcysteine (NAC), they had a lower incidence of early ischemia/reperfusion arrhythmias when compared to control (22). However, no protection against arrhythmic events was observed in cardiomyocytes in which CaMKII methionine residues 281 and 282 were mutated to valine (MMVV) (22). This finding corroborates our previous results showing that methionine 281 and 282 are not the CaMKII oxidation site (8).

### C273S mutation does not affect turnover rate of CaMKIIδ

Mutation of Met281 and Met282 to valine (MMVV mutant) was shown to protect mice and flies against cardiopulmonary pathology induced by oxidative stress (24-26). This provided indirect support for the original report that oxidative stress caused oxidation of Met281/Met282 to the sulfoxide (5). With the recognition that the actual oxidation was a disulfide formation between Cys273 and Cys290, a different explanation was required for protection engendered by the MMVV mutation. We found that the MMVV mutant was degraded four times faster than the wild-type enzyme in HEK293 cells (8). The increased rate of degradation of the MMVV CaMKIIδ could explain the protective effect of the mutation. We considered that the protective effect of the Cys273S mutation might also be due to more rapid turnover of CaMKIIδ. We therefore measured the rates of degradation of the wild-type and mutant proteins (27). Their rates of turnover were identical (Fig. 4A and 4B).

**Figure 4.**
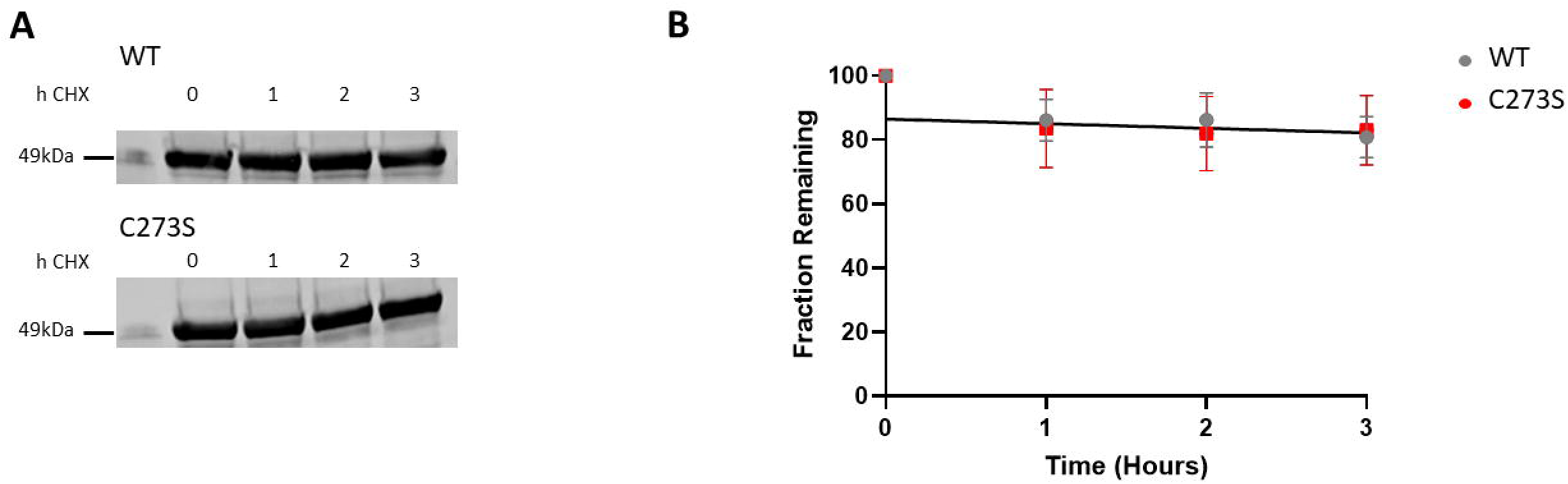
Rate of degradation of wild-type and C273S CaMKIIδ in HEK293T cells. **(A)** HEK293T cells transfected with wild-type or C273S CaMKIIδ were treated with 50 μg/ml cycloheximide (CHX) for 1, 2, or 3 hours. Cells were lysed, and the supernatant was probed with anti-CaMKII antibody. Representative image of n = 3 replicates. **(B)** CaMKIIδ remaining wild-type (grey) and C273S CaMKIIδ (red). The level of CaMKIIδ at 0 h was set to 100%.

Although ROS has been shown to play an important role in heart diseases, no improvement was observed when certain antioxidants were used to prevent or treat heart conditions (28-30). KN62 and KN93 are commonly used as CaMKII inhibitors. They inhibit by preventing Ca^+2^/CAM binding to the enzyme, but they are not able to inhibit CaMKII autonomous activation. They have also been shown to inhibit other calmodulin-dependent kinases (31, 32). Peptide inhibitors designed to prevent CaMKII-substrate binding can inhibit both CaMKII Ca^+2^/CAM dependent and autonomous activation induced by phosphorylation. The same has been shown for the CaMKII-specific ATP-competitive inhibitors (31, 32). It is not clear whether these inhibitors would be efficient in blocking CaMKII autonomous activation induced by oxidation.

A better understanding of the CaMKII activation by oxidation mechanism may facilitate the development of new treatment strategies. It would be useful to have an inhibitor that would only target oxidized CaMKII, since CaMKIIδ has a central role in regulating various cellular processes in the heart.

## Conclusion

We conclude that in the Langendorff *ex vivo* model, CaMKIIδ^C273S^ hearts were protected against ischemia-reperfusion injury and that protection was not due to a change in protein turnover. These results are consistent with our previous finding that under oxidative stress, cysteine 273 is the site of CaMKIIδ oxidation. Further *in vivo* studies will be necessary to determine whether the CaMKIIδ^C273S^ mouse is protected from ischemia/reperfusion injury, as well as the effect on CaMKIIδ downstream phosphorylation targets. While targeting a specific Cys residue is challenging, it is now feasible (33, 34). Thus, pharmacologic prevention of oxidation of Cys273 may be achievable and could be therapeutic in human cardiac disease.

## Supporting information

NIH Publishing Agreement

## Acknowledgment

We thank the Murine Phenotyping Core at NHLBI/NIH for the cardiac physiology tests and the Transgenic Core at NHLBI/NIH for the generation of the CaMKIIδC273S knockin mouse line.

## Data availability

The raw data files have been deposited at our Figshare site, https://doi.org/10.25444/nhlbi.29561903.

## Funding and additional information

This work was supported by the National Institutes of Health NHLBI Intramural Research Program grant ZIA HL000225 to RLL. The content is solely the responsibility of the authors and does not necessarily represent the official views of the National Institutes of Health.

## Author contribution

Nathália Rocco-Machado: Conceptualization, Methodology, Validation, Formal analysis, Investigation, Writing - Original Draft. Junhui Sun: Methodology, Validation, Formal analysis. Audrey Noguchi: Methodology. Danielle A. Springer: Methodology, Formal analysis. Chengyu Liu: Methodology, Validation, Formal analysis. Elizabeth Murphy: Methodology. Rodney L. Levine: Conceptualization, Methodology, Resources, Writing - Review & Editing.

## Graphical Abstract

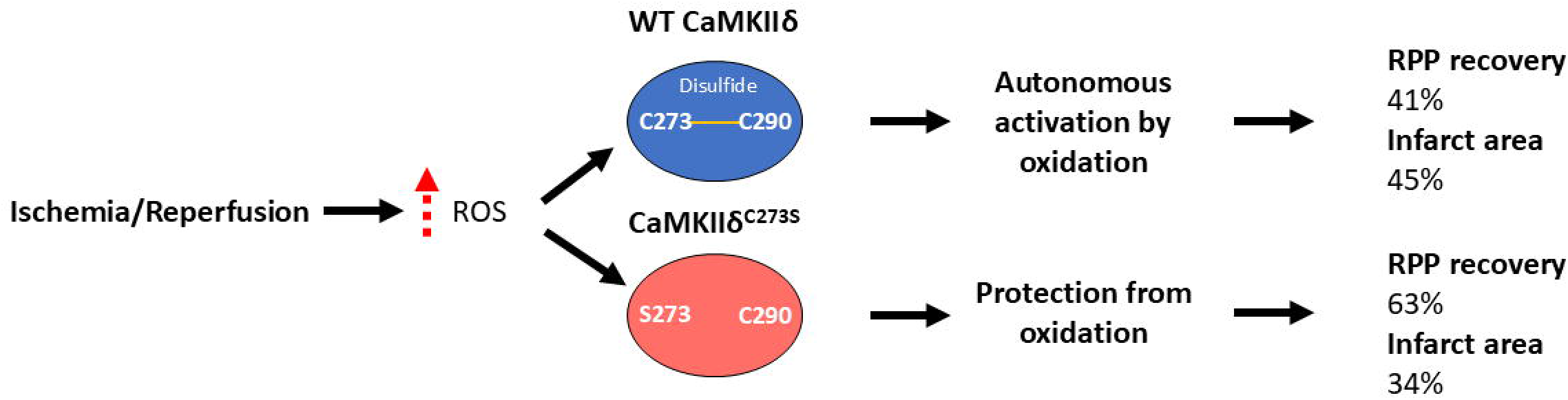

