## Supplemental Figure 1 for "Mutation of a single cysteine in CaMKIIδ protects the heart from ischemia-reperfusion Injury"

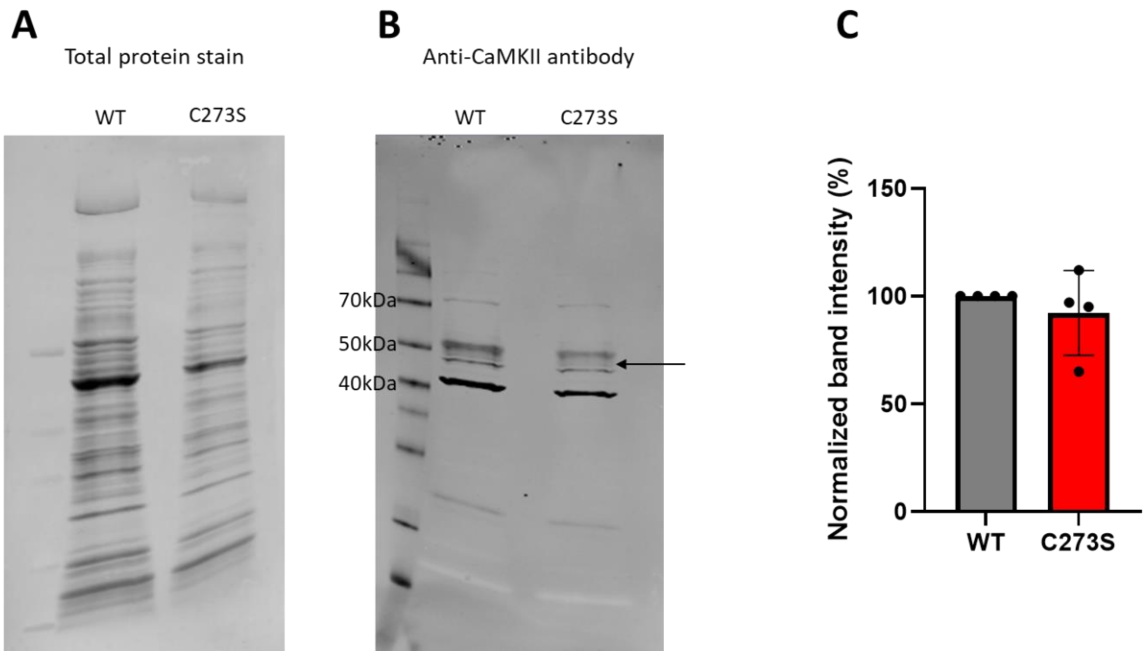


**Supplementary Figure 1. The C273S mutation did not alter CaMKII expression level in the heart. (A)** CaMKIIδC273S and wild-type mice heart lysate supernatant total protein stain. **(B)** CaMKIIδC273S and wild-type mice heart lysate supernatant were probed with anti-CaMKII antibody. Representative image of n = 4 replicates. **(C)** CaMKII band intensity around 50kDa quantification. Protein loading was normalized by total protein staining.

**Raw images**

Experiment 1

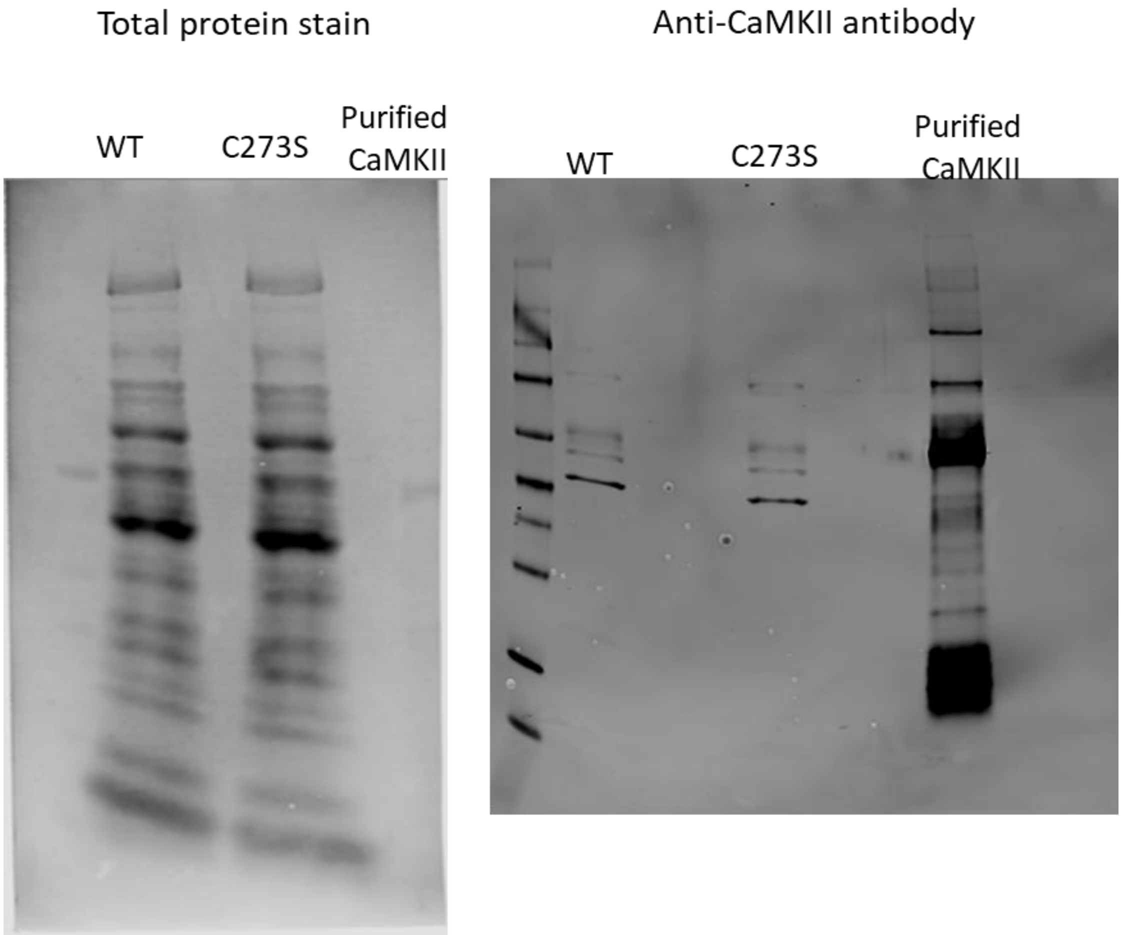

Experiment 2

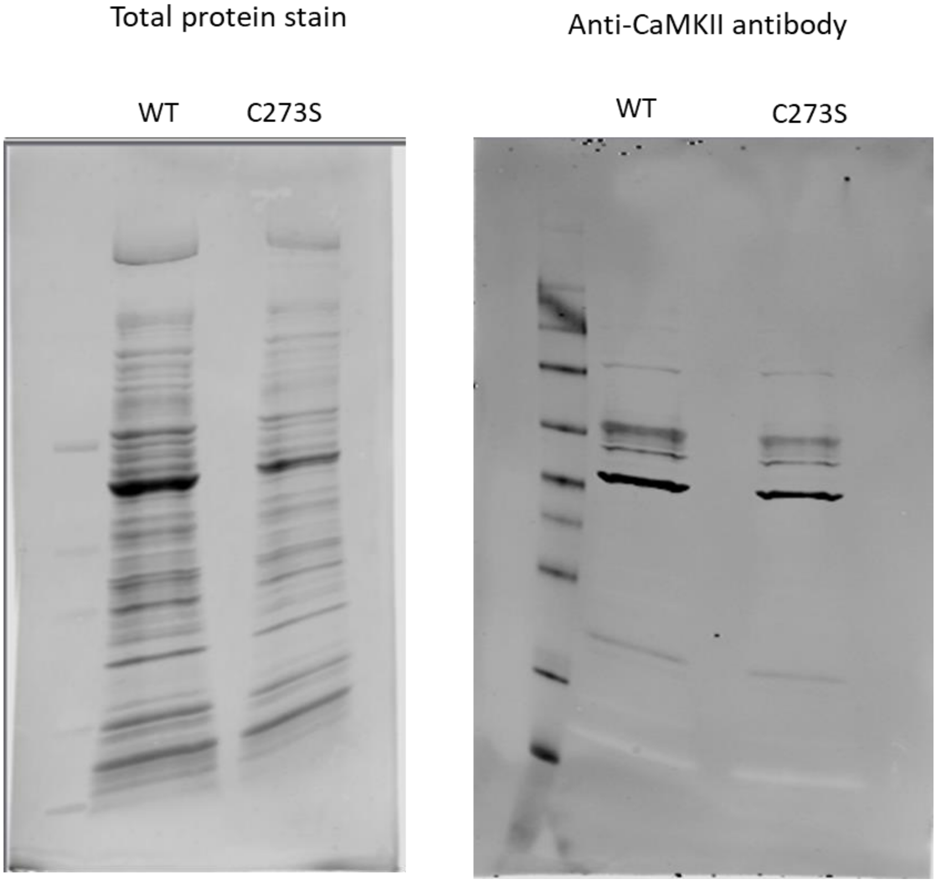

Experiment 3 and 4

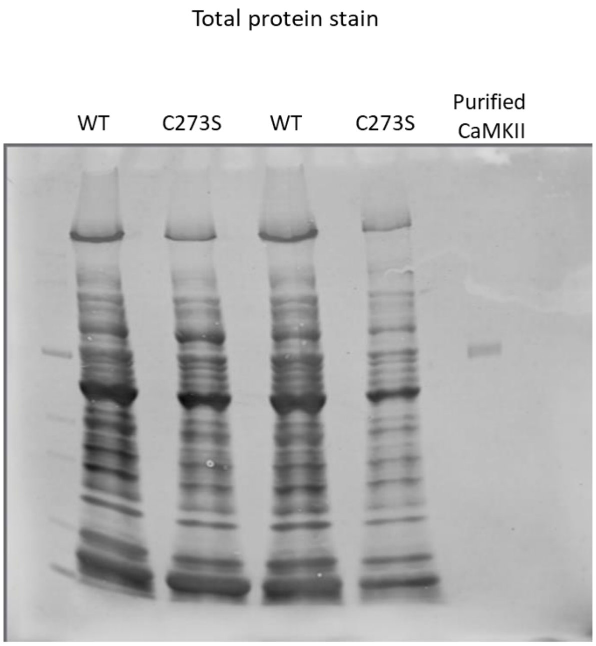


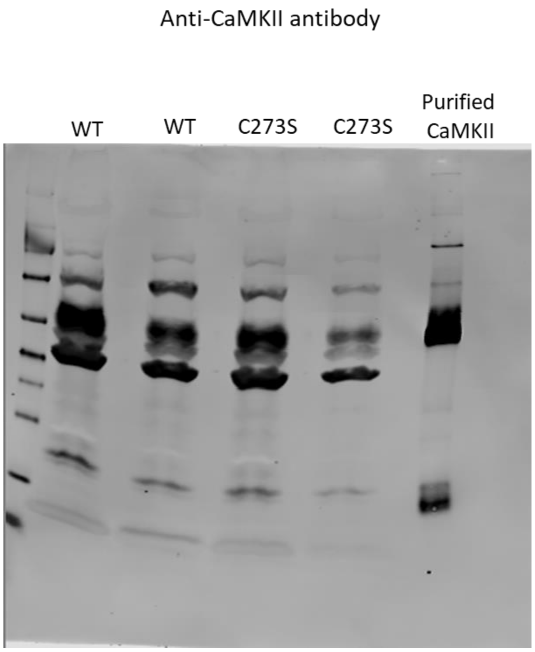
