## Supplementary material for "Mutation of a single cysteine in CaMKIIδ protects the heart from ischemia-reperfusion Injury": NIH Publishing Agreement

### NIH Intramural Research Program and Employee Publishing Agreement & Manuscript Cover Sheet

By signing this Cover Sheet, the Author, on behalf of NIH, agrees to the provisions set out below, which modify and supersede, solely with respect to NIH, any conflicting provisions that are in the Publisher's standard copyright agreement (the "Publisher's Agreement"). If a Publisher's Agreement is attached, execution of this Cover Sheet constitutes an execution of the Publisher's Agreement subject to the provisions and conditions of this Cover Sheet.

1. **Indemnification.** No Indemnification or "hold harmless" obligation is provided by either party.

2. **Governing Law.** This agreement will be governed by the law of the court in which a claim is brought.

3. **Copyright.** Author's contribution to the Work was done as part of the Author's official duties as an NIH employee and is a Work of the United States Government. Therefore, copyright may not be established in the United States. 17 U.S.C. § 105. If Publisher intends to disseminate the Work outside of the U.S., Publisher may secure copyright to the extent authorized under the domestic laws of the relevant country, subject to a paid-up, nonexclusive, irrevocable worldwide license to the United States in such copyrighted work to reproduce, prepare derivative works, distribute copies to the public and perform publicly and display publicly the work, and to permit others to do so.

4. **No Compensation.** No royalty income or other compensation may be accepted for work done as part of official duties. The author may accept for the agency a limited number of reprints or copies of the publication.

5. **NIH Representations.** NIH represents to the Publisher that the Author is the sole author of the Author's contribution to the Work and that NIH is the owner of the rights that are the subject of this agreement; that the Work is an original work and has not previously been published in any form anywhere in the world; that to the best of NIH's knowledge the Work is not a violation of any existing copyright, moral right, database right, or of any right of privacy or other intellectual property, personal, proprietary or statutory right; that where the Author is responsible for obtaining permissions or assisting the Publishers in obtaining permissions for the use of third party material, all relevant permissions and information have been secured; and that the Work contains nothing misleading, obscene, libelous or defamatory or otherwise unlawful. NIH agrees to reasonable instructions or requirements regarding submission procedures or author communications, and reasonable ethics or conflict of interest disclosure requirements unless they conflict with the provisions of this Cover Sheet. Notwithstanding the foregoing, the Author may, consistent with NIH policy, submit a copy of the original work to a public repository.

6. **Disclaimer.** NIH and the Author expressly disclaim any obligation in Publisher's Agreement that is not consistent with the Author's official duties or the NIH mission, described at https://www.nih.gov/about-nih. NIH and the Author do not disclaim obligations to comply with a Publisher's conflict of interest policy so long as, and to the extent that, such policy is consistent with NIH's own conflict of interest policies.

7. **For Peer-Reviewed Papers to be Submitted to PubMed Central.** The Author is a U.S. government employee who must comply with the NIH Public Access Policy, and the Author or NIH will deposit, or have deposited, in NIH's PubMed Central archive, an electronic version of the final, peer-reviewed manuscript upon acceptance for publication, to be made publicly available upon the Official Date of Publication, defined in the NIH Public Access Policy as the date on which the Final Published Article is first made available in final, edited form, whether in print or electronic (i.e., online) format. The Author and NIH agree (notwithstanding Paragraph 3 above) to follow the manuscript deposition procedures of the publisher so long as they are consistent with the NIH Public Access Policy, including making the manuscript publicly available without embargo in PubMed Central.

8. **Modifications.** PubMed Central may tag and modify the work consistent with its customary practices and within the meaning and integrity of the underlying work.

The NIH Deputy Director for Intramural Research approves this publishing agreement for NIH staff from the NIH Intramural Program. NIH Institute and Center Directors approve this publishing agreement for NIH staff from their respective Institutes and Centers outside the NIH Intramural Program. The NIH Principal Deputy Director approves this publishing agreement for NIH staff in the Office of the Director. A single, signed copy of this text is maintained for all works published by NIH employees, and contractors and trainees who are working at the NIH. No additional signatures beyond that of the Author are needed.

Author's name: Nathalia Rocco Machado

Author's Institute, Center, or OD Office: NHLBI

Check if from the NIH Intramural Research Program ☑
Check if Publisher Agreement is attached ☑

Title of Manuscript/Work: Mutation of a single cysteine in CaMKIIδ protectsthe heart from ischemia-reperfusion Injury

Name of publication: bioRxiv preprint

Nathalia Rocco Machado
Author's signature

04/23/2026
Date

NIH OGC May 2025
